# First report on the effective intraperitoneal therapy of insulin-dependent Diabetes mellitus in pet dogs using “Neo-Islets,” aggregates of adipose stem and pancreatic islet cells

**DOI:** 10.1101/666644

**Authors:** Anna Gooch, Ping Zhang, Zhuma Hu, Natasha Loy Son, Nicole Avila, Julie Fischer, Gregory Roberts, Rance Sellon, Christof Westenfelder

## Abstract

We previously reported that allogeneic, intraperitoneally administered “Neo-Islets,” composed of cultured pancreatic islet cells co-aggregated with high numbers of immunoprotective and cytoprotective Adipose-derived Stem Cells, reestablished, through omental engraftment, redifferentiation and splenic and omental up-regulation of Regulatory T-cells, normoglycemia in autoimmune Type-1 Diabetic Non-Obese Diabetic (NOD) mice without the use of immunosuppressive agents or encapsulation devices. Based on these observations, we are currently testing this Neo-Islet technology in an FDA guided Pilot Study (INAD 012-776) in insulin-dependent, spontaneously diabetic pet dogs by the intraperitoneal administration of 2×10e5 Neo-Islets/kilogram body weight to metabolically controlled (blood glucose, triglycerides, thyroid and adrenal functions) animals under sedation and local anesthesia and ultrasound guidance. We report here initial observations on the first 4 Neo-Islet-treated, insulin dependent pet dogs that are now in the intermediate-term follow-up phase of the study (> 6 months post treatment). Current results indicate that in dogs, Neo-Islets appear to engraft, redifferentiate and physiologically produce insulin, and are neither rejected by auto- or allo-immune attacks, as evidenced by (a) an absent IgG response to the allogeneic cells contained in the administered Neo-Islets, and (b) progressively improved glycemic control that achieves up to a 50% reduction in daily insulin needs paralleled by a significant fall in serum glucose levels. This is accomplished without the use of anti-rejection drugs or encapsulation devices. No adverse or serious adverse events related to the Neo-Islet administration have been observed to date. We conclude that this minimally invasive therapy has significant translational relevance to veterinary and clinical Type 1 Diabetes Mellitus by achieving complete and at this point partial glycemic control in two species, i.e., diabetic mice and dogs, respectively.

## Introduction

Diabetes Mellitus is a common endocrine disorder in dogs, and it is estimated that there are currently 700,000 insulin-dependent pet dogs in the US [1–4]. Their care is burdensome and expensive for their owners. As in humans, Type 1 Diabetes Mellitus (T1DM) in dogs is caused by lack of insulin secretion in response to glucose, resulting in hyperglycemia, acid-base and electrolyte disorders, polydipsia, polyuria and weight loss, and is accompanied by a broad spectrum of diabetes-induced end organ and other complications, including blindness due to retinopathy and cataracts, opportunistic infections, neurological and other serious micro- and macro-vascular complications [4–6]. Although dogs were the model in which insulin was originally discovered, and remain a major large animal model for the refinement of diabetic treatments such as pancreas and islet cell transplants, almost no advances in the treatment for diabetic dogs have been made in the last 50 years [7]. A few studies have examined xeno- or allogeneic islet transplantation to reverse or ameliorate diabetes in dogs and have had varying degrees of success. Yet, insulin replacement therapy and blood glucose monitoring remain the only currently available therapy for these animals [7–9]. Due to the challenges of medically managing a diabetic dog, up to 40% of owners regrettably opt to euthanize their dogs within a day of diagnosis rather than treat them [1,10].

While the pathogenic mechanisms of canine T1DM are still incompletely understood, there is evidence that autoimmunity plays a role in approximately 1/3 of cases [3,4,7,10–12]. T1DM occurs with equal frequency in male and female neutered dogs, but as with Non-Obese Diabetic (NOD) mice, at higher frequency in intact females vs. males, suggesting a role for female hormones in the development of the disease in dogs [1,3,13–15]. While T1DM affects both juvenile and adult dogs [1,3,7], it is more commonly seen in adults, generally diagnosed between the ages of 3 and 15 years [3,10]. Some groups have reported isolation of auto-antibodies to proinsulin, GAD65 and IA-2 from the sera of diabetic animals [16,17]. Others, however, have been unable to confirm the presence of such auto-antibodies in the same, previously tested sera, nor in sera from other diabetic dogs [2,14]. On the other hand, several studies have found a genetic association between certain dog leukocyte antigen alleles (DLA) and the development of DM in dogs, similar to that found between HLA alleles and the development of DM in humans [12,18,19]. Despite some controversy as to the immune-mediated destruction of beta cells, all pioneering work on islet and pancreas transplantation for humans was carried out in dogs and clearly demonstrated the need for immunosuppression or immune-isolation, as well as sufficient nutrition/oxygenation and vascularization for islet allo-graft survival [7,20].

Dog survival time post the diagnosis of diabetes is short [1,10]. In one study, median post-diagnosis survival time was found to be only 57 days, due either to pet owners’ unwillingness to care for a diabetic animal, or to the dog suffering from advanced stages of diabetic complications at the time of diagnosis. For dogs surviving beyond the first day after diagnosis, the median survival time was 2 years [1]. These low survival rates, and clear unwillingness of some owners to care for diabetic animals, underscore the need for novel and effective therapeutics that remove much of the burden of diabetes treatment and maintenance from pet owners, and to facilitate the survival of affected dogs.

We previously demonstrated that allogeneic, intraperitoneally administered “Neo-Islets” (NIs), composed of culture expanded islet cells (ICs) co-aggregated with high numbers of immunoprotective and cytoprotective Adipose-derived Stem Cells (ASCs), could reestablish normoglycemia in NOD mice with autoimmune T1DM without the use of encapsulation devices or immunosuppressive agents [21]. Glycemic control was similarly achieved using dog-derived ASCs and ICs in a Streptozotocin (STZ) model of diabetes in NOD/SCID mice [21]. Dose finding studies indicated that a dose of 2×10e5 canine Neo-Islets (cNIs) per kg b.wt. would be sufficient to control blood glucose levels [21].

Based on these studies, we initiated an FDA-CVM guided Pilot Study (INAD 012-776) to assess the (i) safety, (ii) feasibility and (iii) efficacy of allogeneic cNIs in significantly reducing or eliminating the need for exogenous insulin in spontaneously diabetic, autoimmune or insulin resistant, insulin-dependent pet dogs. We further assessed whether the administered NIs elicited an allo-immune response. This Pilot study is currently ongoing at Veterinary Specialty Hospital in San Diego, CA, and at Washington State University in Pullman, WA. Six dogs have been treated and four followed for more than 6 months. We report here on the course of the first four NI treated dogs that have been followed for more than 6 months. The overall rationale of demonstrating that this NI therapy is also effective in a second, larger diabetic mammal, the dog, is the fact that this will further strengthen the justification for the currently planned conduct of a clinical trial in study subjects with T1DM.

## Materials and Methods

### Reagents

Reagents used and their manufacturers are listed as indicated below, except for PCR reagents and primers which are listed in S1 Table.

### Study Design

An FDA guided pilot study (INAD 012-776) conducted with IACUC approval at (a) Washington State University in Pullman, WA (WSU) and (b) the Veterinary Specialty Hospital in San Diego, CA (VSH). Ten Insulin dependent pet dogs are included. Eight dogs have been enrolled according to the criteria in Table 1. Informed consent was obtained from all dog owners prior to enrollment. One owner withdrew her dog prior to treatment, 6 dogs have been treated, and 4 of those (VSH-01, VSH-02, WSU-01 and WSU-02) have been followed for 6 months or longer. Enrolled dogs’ demographics are shown in Table 2 and comorbidities in Table 3. Dogs are screened and followed as shown in S2 Table. Pre-treatment serum samples from the treated dogs were tested for the presence of islet autoantibodies (see below for details), and all dogs were examined for comorbidities. After blood glucose and triglyceride levels were optimally controlled, 2×10e5 allogeneic NIs per kg b.wt. were given i.p.. In all animals, blood Glucose and Fructosamine levels, insulin requirements, body weights, food intake, formation of antibodies to allogeneic NIs, animal activity and the development of adverse events were closely monitored by the PIs and the primary veterinarians for each dog.

**Table 1.** Enrollment Criteria.

|  |  |
| --- | --- |
| Enrollment Criteria | • Insulin dependent, diabetic dog on established insulin and diet regimen<br>• Weight between 5 and 12 kg<br>• Spayed, if female |
| Exclusion Criteria | • History of malignancy<br>• Significant illness unrelated to the diabetic state such that the PI believes the dog to be a poor candidate<br>• Significantly advanced age such that the PI believes the dog to be a poor candidate<br>• Contraindication to general anesthesia<br>• Participation in another, ongoing clinical trial |
| Specific Enrollment Criteria Related to Diabetes Mellitus in Dogs | Presence of<br>• End-organ damage<br>• Cataracts<br>• Neuropathy<br>• Renal disease<br>• History of pancreatitis |

**Table 2:** Study Subject Demographics.

| Subject # | Treated | Gender | Breed | Weight at enrollment (kg) | Approx. Age at enrollment (yrs) | Duration of DM (yrs) | insulin dose/day, regimen at enrollment or treatment (later) |
| --- | --- | --- | --- | --- | --- | --- | --- |
| VSH-01 | yes | M | French Bulldog | 12.3 | 9 | 2.5 | 4.5-5.5 U Novalin BID |
| VSH-02 | yes | M | Bichon Mix | 6.9 | 7 | 0.5 | 5 U Vetsulin BID |
| VSH-03 | yes | F | Bichon Poodle Mix | 5.8 | 11 | 2 | 7.5 U Vetsulin BID |
| VSH-04 | yes | M | Chihuahua Mix | 7.3 | 12 | 1 | 9 U Vetsulin BID |
| VSH-05 | stabilizing | M | Poodle | 5.3 | 7 | 0.2 | 7 U Vetsulin BID |
| VSH-06 | withdrawn | M | Chihuahua Mix | 7 | 7 | 1 | 6 U Vetsulin BID |
| WSU-01 | yes | F | American Eskimo | 11 | 1.8 | 0.8 | 2.5 U Vetsulin BID |
| WSU-02 | yes | M | Chihuahua Mix | 7 | 1 | 0.5 | 4 U Vetsulin BID |

Abbreviations: U = Units, WSU = Washington State University, VSH = Veterinary Specialty Hospital, BID = twice per day. Insulin is listed as per dose. If given BID, double for the daily dose.

**Table 3:** Clinical data on treated dogs.

|  |  |
| --- | --- |
| **VSH-01**<br>• Male, neutered (DOB 8/8/08)<br>• No history of pancreatitis, but evidence on screening US of suspected chronic or previous pancreatitis<br>• Cataracts; moderate adrenomegaly, mild renal degenerative changes | **VSH-02**<br>• Male, neutered (DOB 3/28/11)<br>• No history of pancreatitis, but evidence on screening US of suspected chronic or previous pancreatitis<br>• Hypothyroid (0.1 mg Synthroid BID), diagnosed through screening; mild renal degenerative changes |
| **VSH-03**<br>• Female, spayed (DOB 6/19/06)<br>• No history or evidence of pancreatitis<br>• Cataracts, hepatomegaly, elevated ALP, hyperlipidemia (150 mg Gemfibrozil BID) | **VSH-04**<br>• Male, neutered (DOB 10/10/05)<br>• No history or evidence of pancreatitis<br>• Cataracts, hepatomegaly, elevated ALP + ALT, hyperlipidemia (150 mg Gemfibrozil BID) |
| **WSU-01**<br>• Female, spayed (DOB 6/15/16)<br>• No history nor evidence of pancreatitis<br>• Hypercholesterolemia<br>• History of liver disease | **WSU-02**<br>• Male, neutered (DOB 4/15/17)<br>• No history nor evidence of pancreatitis |

## Blood glucose levels

Blood glucose levels were assessed by the owner at least twice daily (morning and evening) using either an *AlphaTrak* or *FreeStyle Libre* glucometer prior to insulin administration. Insulin (Vetsulin or Novalin) was administered BID based on blood glucose readings, and the doses given are recorded.

## Cells

### Cell donor information

All cells, Adipose-derived Stem Cells (ASCs) and Islet Cells (ICs), for the study were obtained through an NIH sharing agreement from 27 dogs being euthanized under a University of Utah IACUC approved protocol. This study involved no use of test agents, but did involve the use of a surgically implanted pacemaker in some dogs to induce heart failure, as well as the use of resynchronization therapy via the pacemaker as an experimental treatment. Dogs were up-to-date on vaccinations at the time of euthanasia.

### slets and ASCs

Islets and ASCs were isolated and cultured from dogs as previously reported [21–26], and as described in detail in the Supporting Information section of our previous publication [21]. Prior to NI formation, cultured ASCs were characterized for their ability to undergo trilineage (adipoosteo-, chondrogenic) differentiation as described [21] and for surface marker expression of CD90, CD44, CD34, CD45 and DLA-DR as in our previous publication [27], and using the following antibodies: Phycoerythrin (PE)-labeled, monoclonal, rat anti-dog CD90 IgG2b, and Isotype (Invitrogen, 12-5900 and 12-4031-83); Allophycocyanin (APC)-labeled, monoclonal, mouse anti-dog CD44 IgG1; and Isotype (R&D, FAB5449A and IC002A); PE-labeled, monoclonal, mouse anti-dog CD34 IgG1, and Isotype (BD Biosciences, 559369 and 554680); R- PE-labeled, monoclonal, rat anti-dog CD45 IgG2b, and Isotype (BioRad MCA1042PE and PA5- 33195), PE-labeled, monoclonal, mouse anti-human HLA-DR IgG2a (cross-reacts with dog), and Isotype (BD Biosciences 555812 and 555574). All antibodies were used at the concentrations recommended by their respective manufacturers.

## Cell banking

Passage 0 (P0) cultured islet cells and P2 ASCs were suspended in CryoStor CS10 (BioLife Solutions, 210102) and banked, frozen in liquid nitrogen (−140°C) until ready for final expansion and NI formation. Prior to freezing, cells were release tested for viability, sterility, endotoxin, mycoplasma, expression of various genes involved in immune modulation, cell survival and angiogenesis, and dog-specific adventitious agents.

## rtPCR

rtPCR was carried out as described in our previous publication [21] using the reagents and primers listed in S1 Table. In brief, RQ was calculated through normalization to internal (deltaCT; beta actin and beta 2 microglobulin) and external controls (delta-deltaCT; parent cells), both accomplished using the ABS 7500 Real Time PCR System and software. Results are presented as log10(RQ) ± log10(RQmin and RQmax). Differences between expression levels greater than log10(RQ) 2 or log10(RQ) -2 were considered significant.

## Final Product (NI) formation and storage

NIs were formed in ultra-low adherent 10-layer Cell Stacks (Corning, custom made product) from freshly cultured banked ICs and ASCs using 70×10e6 cells per layer and 140 ml of DMEM (Gibco 11885-084) + 10% dog serum (Golden West) as described in our previous publication [21]. NIs were harvested and resuspended in 50 to 100 ml of sterile Plasmalyte A (Baxter, 2B2543) + 2% HEPES (Gibco), pH 7.4 at a concentration of 2×10e7 clustered cells/ml, and placed in a sterile 100 ml syringe (Wilburn Medical, WUSA/100). The final product was release tested (viability, sterility, endotoxin, mycoplasma, gram stain, gene expression of insulin (INS), glucagon (GCG), somatostatin (SST), pancreatic polypeptide (PPY), pancreatic and duodenal homeobox 1 (PDX1), vascular endothelial growth factor A (VEGFa), stromal cell derived factor 1 (CXCL12), and stored and transported to the study site at 4°C for administration within 48 hrs of packaging.

## Cell bank and final product release testing

Cell viability was assessed using Fluorescein diacetate (Sigma, F7378) and Propidium Iodide (Life Technologies P3566) as per the manufacturers’ instructions. Sterility was assessed as described in 21 *CFR 610*.*12*, using Tryptic Soy Broth (Sigma, 22092) and Fluid Thioglycollate Medium (Sigma, T9032) and following the manufacturer’s instructions. Endotoxin levels were determined using the Charles River Endosafe Nexgen PTS system and reagents (Charles River) following the manufacturer’s instructions. Possible Mycoplasma contamination was assessed using the MycoAlert PLUS Mycoplasma Detection Kit (Lonza, LT07-701 and LT07-518) per the manufacturer’s instructions. Samples of banked cells from each donor dog were sent to Zoologix for relevant adventitious agent testing. Prior to NI formation, ASCs were assessed for expression of genes involved in immune modulation, cell survival and angiogenesis. ICs were analyzed for expression of islet hormone associated genes. Gram Staining was conducted by standard methods using a kit (Sigma, 77730-1KT-F). Cell and NI release criteria are listed in Table 4.

**Table 4:** Release criteria for cells and Neo-Islets.

|  | Viability | Sterility | Endotoxin | Mycoplasma | Adventitious Agents | Gene expression (rtPCR) | extracellular marker (FACS) |
| --- | --- | --- | --- | --- | --- | --- | --- |
| **ASCs** | ≥ 70% | No growth after 14 days | < 5.0 EU/ml | B/A <1.2 | negative for all tested | Significant increase in Ido 1 expression Upon overnight culture with INF-γ | ≥ 90% CD90, CD44; ≤ 4% CD34, CD45, DLA-DR |
| **Islet Cells** | ≥ 70% | No growth after 14 days | < 5.0 EU/ml | B/A <1.2 | negative for all tested | + Ins | NA |
| **Neo-Islets** | ≥ 70% | negative gram stain; no growth after 14 days | < 5.0 EU/ml | B/A <1.2 | NA | + Ins | NA |

Adventitious Agents tested:
Neorickettsia risticii, Mycoplasma haemocanis, Mycoplasma canis, Bartonella henselae, B. bacilliformis, B. clarridgeiae, B. elizabethae, B. quintana and B. vinsonii subsp. Berkhoffii, Brucella abortus, B. microti, B. melitensis, B. pinnipedialis, B. suis, B. canis, B. ovis and B. neotomae, Neorickettsia helmintheca, Cryptococcus neoformans, Influenza A H5N1, H5N2, H1N1, H2N2, H3N8, H4N6, H7N7, H8N4 and H9N2, canine herpesvirus, Toxoplasma gondii

## Testing of treated dogs’ sera for antibodies to Islet Cells and ASCs

Prior to and at ∼6 weeks post NI treatment, 1 ml of serum was collected from each study dog. 5×10e4 banked allogeneic ASCs, and (separately) 5×10e4 banked allogeneic ICs were incubated for 30 min at room temperature with 100 uL serum each. Cells were then centrifuged at 600 x g for 5 min; the supernatant was removed and the cell pellets resuspended in 200 uL Phosphate Buffered Saline (PBS, Sigma, 11666789001) +1% Fetal Bovine Serum (FBS, GEHealthcare, SH30910.03). Cells were then incubated at room temperature for 30 min with a 1:200 dilution of FITC-labeled, polyclonal rabbit anti-dog IgG or Isotype control (Jackson ImmunoResearch; 304- 095-003, lot #13971 and 011-090-003, lot #127025 respectively). They were then centrifuged, washed 1x with PBS + 1% FBS; resuspended in 400 uL PBS + 1% FBS, and analyzed by FACS as previously described [21].

## Enzyme Linked Immunosorbent Assays (ELISAs)

ELISAs for GAD65 and IA2 autoantibodies were kindly carried out by Dr. Boris Fehse, University of Hamburg, Germany, using GAD65 and IA2 ELISA kits (both from Euroimmun, Luebeck, Germany, EA 1022-9601 G and EA 1023-9601 G, respectively) and the same serum samples were analyzed for allo-antibody levels. The GAD65 kit has previously been shown to cross-react with dog antibodies [28].

## NI Administration

NIs were administered to dogs by the site’s veterinary ultrasonographer as follows: an i.v. catheter was placed, and the dog was sedated with dexmedetomidine (5 mcg/kg) and butorphanol (0.1-0.2 mg/kg). Abdominal fur was shaved, and skin was prepared for aseptic injection. NIs were warmed to room temperature, and an 18 gauge, sterile cannula was placed, after local anesthesia was administered, through the linea alba, ∼ 3-4 cm cranial to the umbilicus, under ultrasound guidance to intraperitoneally infuse normal saline test solution, then the suspended NIs over a 2-3 minute period. Once the administrations were completed, the cannula was withdrawn. Ultrasound imaging was carried out to check for abdominal bleeding. Sedation was reversed with Antisedan, and the dog was monitored until determined stable by the PI and veterinary staff. Dogs returned home with their owners the same day, once the PI determined that they were stable and ready. Owners were advised that the dogs’ serum glucose levels should be kept ≤ 210 mg/dL to protect the graft cells and facilitate redifferentiation into insulin producing cells.

## Follow-up Schedule

### Month 1

Dog owners check and record their dog’s blood glucose levels every 12 hours; record food and water intakes and weights once per week on supplied forms. Dog owners administer insulin to dogs as needed and as instructed by the PI. Dogs are brought in for a physical examination and laboratory studies as indicated in S2 Table.

### Months 2 - 12

Dog owners continue to check and record their dog’s blood glucose levels twice per day; record dogs’ daily food and water intakes and weights every other week on supplied forms and electronically through month 6, and then as deemed necessary by the PI. Dog owners are responsible for the continued administration of insulin to dogs as needed and as instructed by the PI. Dogs are brought in for a physical examination once per month post treatment through month 6, and once per quarter in months 6 - 12. Fructosamine levels, a chemistry panel and urinalysis are obtained at the 3rd and 6th month visits. At the 6 months visit, a CBC, Chemistry and electrolyte panel, HbA1c, and urine are collected. At each visit, potential changes in the degree of preexisting end-organ damage are carefully documented.

### Years 1-3

Dog owners are responsible for continued checking of blood glucose levels and administering insulin as deemed necessary by the PI. Dogs are brought in for a physical examination once per quarter through the 36th month post treatment, to assess changes in the degree of documented end-organ damage. Fructosamine levels, HbA1c and chemistry panels are checked at each visit. Once per year, a CBC and a urinalysis are obtained.

## Statistical Analysis

Unless otherwise indicated, Data are expressed as Mean ± SEM or Mean ± 95% confidence interval, as indicated. Primary data were collected using Excel (Microsoft, Redmond, WA), and statistical analyses were carried our using Prism (GraphPad, San Diego, CA). Two tailed T-tests were used to assess differences between data means. A *P* value of < 0.05 was considered significant.

## Results

### Study Design

This FDA-CVM guided Pilot study to determine the safety, feasibility and preliminary efficacy of cNIs in eliminating or significantly reducing a diabetic dog’s need for insulin will be carried out in 10, stable, insulin-dependent, diabetic pet dogs of either gender as described in Methods. Accordingly, the primary endpoints are safety and feasibility, as well as a demonstrated lack of immune response to the administered cells as evidenced by either sustained euglycemia or improvement in glucose control and/or reduced need for insulin after NI administration; and the absence of IgG antibodies directed at the cells that compose the allogeneic NIs in sera of treated dogs as assessed by FACS at ≥ 1 month post transplantation. The secondary endpoint is reduction or elimination of need for insulin in treated dogs. The tertiary endpoint is lack of development or progression of end-organ damage in treated dogs. Adverse and Severe Adverse Events are recorded and reported over the duration of the study.

Dogs are enrolled according to the criteria in Table 1, treated with NIs when serum glucose and serum lipids are stable and controlled to within normal ranges, and they are monitored for adverse events, and serum glucose levels and insulin needs are recorded according to the schedule in S2 Table, and as indicated in Methods. Additionally, owners check, record and report their dogs’ blood glucose levels and insulin needs twice per day, and monitor their dogs’ food and water and food intake, activity and weight.

At this point, 8 eligible dogs have been enrolled for treatment with allogeneic NIs. The dogs’ pretreatment demographics and comorbidities are summarized in Tables 2 and 3. Of these dogs, one dog’s owner withdrew him from the study prior to dosing, two are being treated for hypertriglyceridemia (Gemfibrozil, 150 mg BID, and dietary restriction) and have been treated but are not yet in the intermediate term follow-up phase of the study, while four have been treated with NIs and (VSH-01, VSH-02, WSU-01 and WSU-02) have been followed for more than 6 months. VSH-01 was a 9 year old (at dosing), male, 12 kg French bulldog who had been on insulin for approximately 2.5 years at the time of dosing. VSH-02 was a 7 year old, hypothyroid, male, 7 kg Bichon mix who had been diabetic for approximately 6 months at the time of dosing. WSU-01 was a 2 year old, 11 kg, female American Eskimo dog who had been on insulin for approximately 9 months prior to treatment. WSU-02 was a 1 year old, 7 kg Chihuahua mix who had been on insulin for approximately 6 months prior to dosing.

Therapeutic Doses of NIs for all treated dogs were prepared, release-tested, packaged and administered as described in Methods. All cells and final products met the release criteria listed in Table 4. Prior to administration, NIs were characterized by rtPCR for gene expression of INS, GCG, SST, PPY, PDX-1, NKX6-1, VEGFA, and CXCL12. As shown in Fig 1, while PDX-1 was no longer expressed in NIs given to any dog, NIs used for treatment were shown to transcribe islet hormone genes for INS and GCG, albeit at significantly reduced levels compared to P0 cultured islet cells. NIs also expressed genes associated with ASCs cytoprotective, angiogenic and immune modulatory activities [29–34] (Fig1).

**Figure 1.**
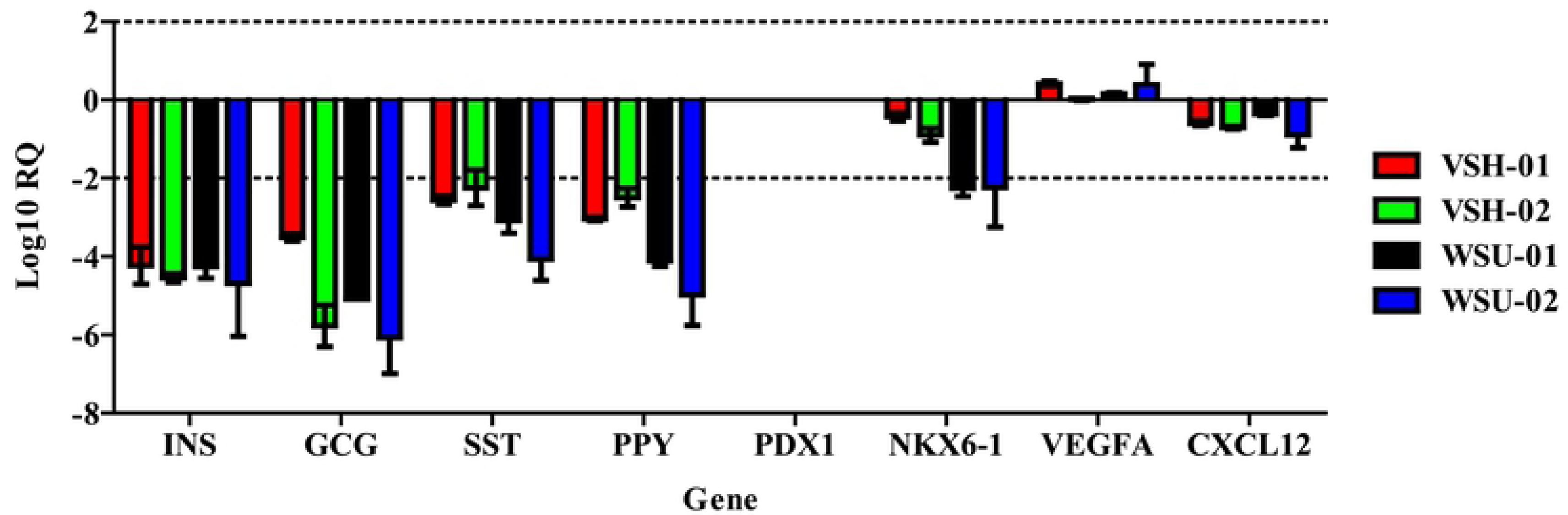
Gene expression profiles (Log10RQ) of the NIs administered to study subjects. All gene expression levels were normalized to those of the P0 IC banks from which the IC portion of the NIs were derived. Data are expressed as mean with 95% Confidence Interval, and reactions were carried out in duplicate. INS, GCG, SST, PPY, NKX6-1, VEGF, and CXCL12 are all expressed in the NIs administered to each dog. PDX-1 is no longer expressed.

## Intermediate term blood glucose and insulin requirements

As shown in Fig 2, blood glucose levels and insulin requirements for VSH-01, VSH-02, and WSU-02 were significantly reduced (P < 0.05) ≥ 6 months post treatment compared to baseline. Although WSU-01’s mean monthly blood glucose levels 12 months post treatment were also significantly reduced compared to those at baseline (P = 0.0015), her daily insulin needs remained unchanged (Fig 2).

**Figure 2.**
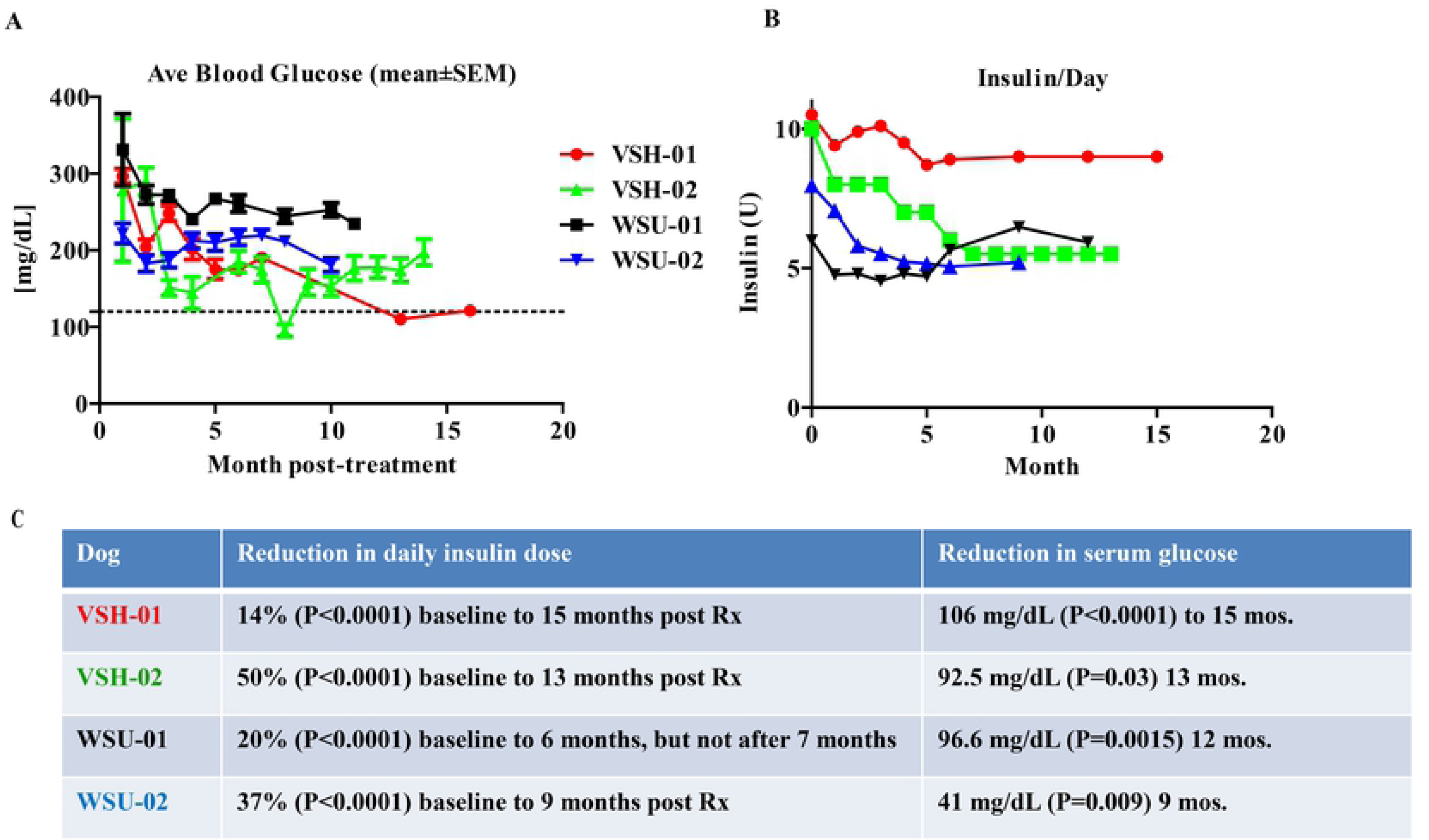
Serum glucose levels and insulin needs over time. (A) Serum glucose levels of study dogs, as assessed and reported by owners, prior to treatment (0 months) and over the study period. As dogs were treated at different times, they are not all currently at the same post-treatment time, thus the reported follow-up period for each is different. (B) Insulin needs for the same time frame. Glucose and insulin are reported at least 2x per day. All glucose level values are averaged for the pretreatment period and for each month post-treatment. Units of insulin administered per day are calculated and averaged for each month. (C) Percent reduction in daily insulin dose at the current follow-up point from baseline and average mg/dL reduction in serum glucose from baseline along with statistical significance (P values) for each dog are shown. With the exception of WSU-01, all treated dogs currently show both a sustained reduction in serum glucose and need for insulin. WSU-01 showed sustained reduction in serum glucose for her entire 12 month follow-up period, but only sustained a 20% reduction in her need for insulin over 6 months post-treatment. After 7 months, her need for insulin rebounded to her pretreatment levels and remains there now.

## Type of diabetes

As part of the study, serum from dogs was obtained prior to and after NI dosing, and tested by FACS for antibodies to the administered cells, using pre-dosing serum and isotypes as negative controls. Pre-treatment testing of sera served not only the purpose of giving a baseline level of anti-ASC or anti-IC antibodies to compare to post-treatment levels, but also as an indication of whether the dogs’ diabetes was auto-immune in nature, as indicated by the presence of anti-Islet Cell antibodies prior to treatment. By this criterion, three of the four tested dogs, VSH-02, WSU- 01 and WSU-02, appear to have autoimmune diabetes (see Fig 3A and B), while VSH-01 may have an “insulin resistance” form of Insulin-dependent DM.

**Figure 3.**
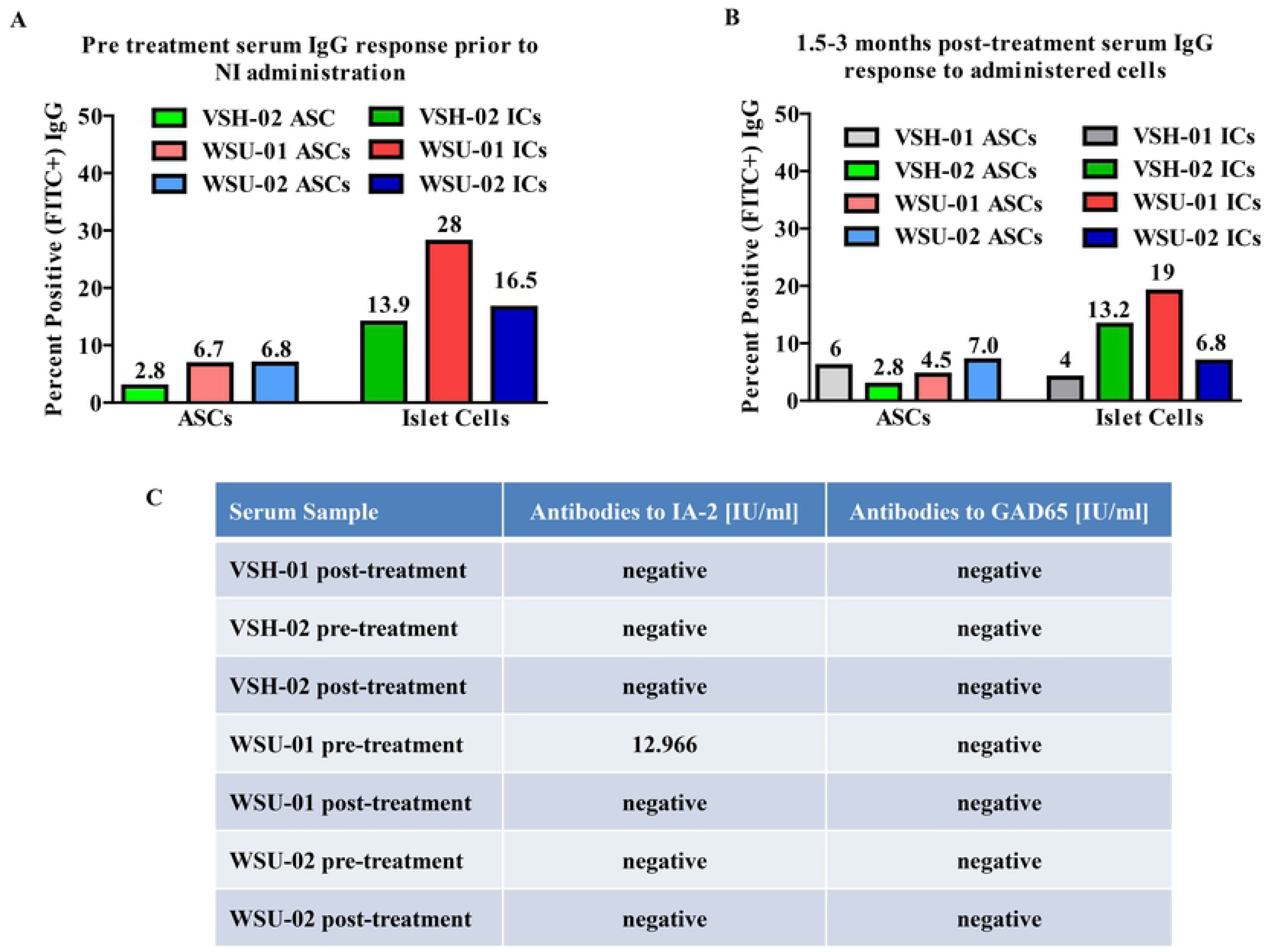
Antigenic responses to NI treatment, and pre-existing presence of auto-islet antibodies. Anti ASC and anti-IC IgG responses as assessed by FACS in sera of the study dogs before (A) and after (B) treatment with allogeneic NIs. Shown are percentages of FITC-labeled anti-dog IgG antibody. Sera were collected from dogs before (A) and 1.5 to 3 months after (B) NI administration. The percent of positive cells is indicated above each column. Dogs do not show increased IgG responses to either ASCs or ICs after NI administration, indicating there is no additional allo-immune response by the recipients to either cell type. Three dogs show pre-existing antibodies to ICs, prior to treatment with NIs, suggesting they have an autoimmune form of diabetes. (C) Results of ELISA testing of sera from the treated dogs for specific anti-islet antigens, IA2 and GAD65, indicate that none of the dogs have antibodies to GAD65 antigen, but that WSU-01’s serum contained antibodies to IA2 prior to, but not after treatment. Samples were run in duplicate. Note: a pre-treatment serum sample was not available for VSH-01.

Sera were similarly tested for the presence of antibodies to two common human autoimmune antigens that have been associated with canine T1DM, IA2 and GAD 65 [16,17]. Only pretreatment serum from WSU-01 showed reactivity to IA2. No other sera showed the presence of antibodies to either (Fig 3C).

## Neither auto- nor allo-immune rejection is observed

Even though at least 3 of the treated dogs (VSH-02, WSU-01 and WSU-02) appear to have autoimmune diabetes (Type 1) as indicated by the presence of anti-islet cell IgG in their sera (see Fig 3), and even though the NIs used to treat them came from unrelated donors, none of the dogs appear to have rejected the NIs as indicated by the following. First, responsive dogs show continued improved blood sugars and lowered insulin requirements (see Fig 2). Second, no allo-rejection antibodies to the NIs are found in the treated dogs’ sera after implantation (Fig 3).

## NI therapy appears to be safe and well tolerated in dogs

In addition to regular blood glucose and weight monitoring, dogs enrolled in this study are being followed closely over the entire 3 year follow-up period with physical examinations and laboratory tests in order to detect signs of adverse events or changes in end organ damage in association with Neo-Islet therapy (see S2 Table).

Despite the fact that several of the currently studied dogs are of advanced age and have multiple comorbidities (see Tables 2 and 3), no adverse events attributable to the NI therapy have been observed to date. Specifically, none of the 6 treated dogs have developed adverse events such as oncogenic transformation of transplanted cells, hematological changes, deterioration in organ function, lack to thrive, etc. While at this point in the study we cannot rule out with certainty the possibility that adverse events can eventually occur, data in the nearly 2 years since the first dogs were treated would indicate that Neo-Islets are safe and well tolerated.

## Discussion

Intermediate term results from the current study thus far demonstrate that allogeneic NI therapy, as currently dosed, (i) is effective in improving glycemic control while durably reducing the need for insulin; (ii) it does so without eliciting an immune response, even in dogs with autoimmune diabetes; and (iii) is feasible and safe. The observed decrease in post-treatment compared to pre-treatment levels of anti-Islet Cell antibodies in two dogs (Fig 3A) may reflect the known inhibitory actions of ASCs on B cells [35]. However, this potentially significant effect must be confirmed in additional studies. While the documented reduction in total insulin requirement occurs only gradually as transplanted ICs re-differentiate into insulin producing cells, this response does taper off subsequently, most importantly, it does not, in most cases, increase again as is seen in failing traditional intrahepatic islet cell transplants (Fig 2) [36]. The data so far indicate that the allogeneic NI grafts are stable and functioning long term, and are not being rejected, which directly demonstrates that this novel form of therapy does not require the life-long use of potentially toxic antirejection drugs. In other words, the allogeneic ASC component of the administered NIs appears to provide through its immune-modulating activity [35] both robust auto- and allo-immune isolation of the cells that make up NIs, and this without the need for often failing encapsulation devices. For example, the use of such a subcutaneously implanted encapsulation device in a clinical trial has proved problematic, as it elicited an inflammatory fibrotic, foreign body type response that resulted in the death of the encapsulated insulin producing cells and thus failure so far of this mode of T1DM therapy [37].

Since the treated dogs are pets and the study is ongoing, the exact engraftment site of the i.p. administered NIs has not been histologically confirmed, although we believe that the main engraftment site is the omentum as we clearly demonstrated in our mouse studies (21). The fact that none of the NI treated dogs developed hypoglycemic episodes furthermore illustrates that insulin secretion by administered NIs remains physiological and occurs into the portal system of the liver, which is physiological. Late post treatment glucose tolerance tests with simultaneous monitoring of canine insulin and C-Peptide release will be conducted in all study dogs.

There are several possible explanations for the incomplete normalization of blood glucose levels and failure to achieve complete insulin independence. These incomplete responses may be related to an inadequate NI dose, the potential need for a second dose, as is routinely done in human islet transplants [36] and as we demonstrated to be effective in diabetic mice, and potentially suboptimal omental uptake and engraftment of NIs. In addition, the need to keep the dog post NI infusion for at least 24 hrs either in a prone or supine position is important since both of these positions facilitate the omental engraftment of administered NIs, while the assumption of an upright position will lead to the translocation of the transplants to the dog’s pelvis, a location that prevents their engraftment in the omentum and failure to function as intended. The current technology exploits the omentum’s ability to both release cells and to take up cell aggregates such as NIs via its milky spots, combined with its excellent arterial blood supply for oxygenation of and glucose sensing by engrafted NIs. And importantly, the omentum’s venous drainage facilitates the physiological delivery of secreted insulin and other islet hormones directly into the portal system of the liver, i.e., a route that is identical to that of the pancreatic veins. Since the liver inactivates up to 50% of received insulin, the post hepatic concentrations of insulin that other insulin-sensitive tissues are exposed to is significantly lower than those insulin levels that are generated by the s.c. injection of insulin that, particularly in higher doses may have adverse systemic effects [38,39].

Finally, the ability to generate more than 50 therapeutic NI doses from a single cadaveric pancreas and MSC donor will significantly improve the availability of this therapy for diabetic dogs and assist their owners with the care of their pets. The cost savings over time, once the NI therapy has been further optimized to durably make diabetic dogs insulin independent are predicted to be significant.

In conclusion, completion of the current study with the remaining dogs will include permitted protocol modifications that we expect to augment the therapeutic efficacy of the so far utilized NI treatment protocol. Nevertheless, we posit that the here presented observations not only provide evidence in support of our hypothesis that NIs when given i.p. engraft in the omentum where they redifferentiate and create a new endocrine pancreas that leads to the establishment of euglycemia and insulin-independence. The proof of principle, i.e., the demonstration that this NI therapy is also effective in a second, larger diabetic mammal, the dog, is definitively significant as this will further strengthen the justification for the currently planned conduct of a optimally designed clinical trial in study subjects with T1DM.

## Acknowledgements

We thank Valorie Wiss at Washington State University for her excellent work as study coordinator. Some of the here-presented data were shown in poster format, and in oral presentations at the 78^th^ and 79^th^ American Diabetes Association Annual Sessions.

## Supporting information

**S1 Table. PCR reagents used and their sources.**

**S2 Table. Follow-up testing schedule.**

